# Kinetochore dynein is sufficient to biorient chromosomes and remodel the outer kinetochore

**DOI:** 10.1101/2023.03.23.534015

**Authors:** Bram Prevo, Dhanya K. Cheerambathur, William C. Earnshaw, Arshad Desai

## Abstract

Multiple microtubule-directed activities concentrate on chromosomes during mitosis to ensure their accurate distribution to daughter cells. These activities include couplers and dynamics regulators localized at the kinetochore, the specialized microtubule interface built on centromeric chromatin, as well as motor proteins recruited to kinetochores and to mitotic chromatin. Here, we describe an *in vivo* reconstruction approach in which the effect of removing the major microtubule-directed activities on mitotic chromosomes is compared to the selective presence of individual activities. This approach revealed that the kinetochore dynein module, comprised of the minus end-directed motor cytoplasmic dynein and its kinetochore-specific adapters, is sufficient to biorient chromosomes and to remodel outer kinetochore composition following microtubule attachment; by contrast, the kinetochore dynein module is unable to support chromosome congression. The chromosome-autonomous action of kinetochore dynein, in the absence of the other major microtubule-directed factors on chromosomes, rotates and orients a substantial proportion of chromosomes such that their sister chromatids attach to opposite spindle poles. In tight coupling with orientation, the kinetochore dynein module drives removal of outermost kinetochore components, including the dynein motor itself and spindle checkpoint activators. The removal is independent of the other major microtubule-directed activities and kinetochore-localized protein phosphatase 1, suggesting that it is intrinsic to the kinetochore dynein module. These observations indicate that the kinetochore dynein module has the ability coordinate chromosome biorientation with attachment state-sensitive remodeling of the outer kinetochore that facilitates cell cycle progression.

## RESULTS & DISCUSSION

During mitosis, three major force generators are implicated in the alignment and biorientation of chromosomes on the spindle: two types of force-generating motor proteins, cytoplasmic dynein and chromokinesins, and the indirect force-generating Ndc80 module, which couples to dynamic microtubules to harness their polymerization dynamics^1–3^ (**Fig. 1A**). The Ndc80 module in metazoans is comprised of the microtubule-binding Ndc80 and Ska complexes, whose cooperation is important for ordered transitions in end-coupled kinetochore-microtubule attachment stability that ensure accurate segregation ^4–9^. As Ska complex recruitment and actions at the kinetochore depend on Ndc80 ^5, 6, 9, 10^, we refer to their coordinated action as that of the Ndc80 module. A number of microtubule dynamics regulators, including kinesin-13 depolymerases, CLASPs, XMAP215 family proteins, and EB family plus-end tracking proteins, are also concentrated at the kinetochore-spindle microtubule interface and are important for proper chromosome segregation ^2, 11^. However, many of these factors act globally on microtubules, which complicates analysis of their specific functions at kinetochores.

**Figure 1.**
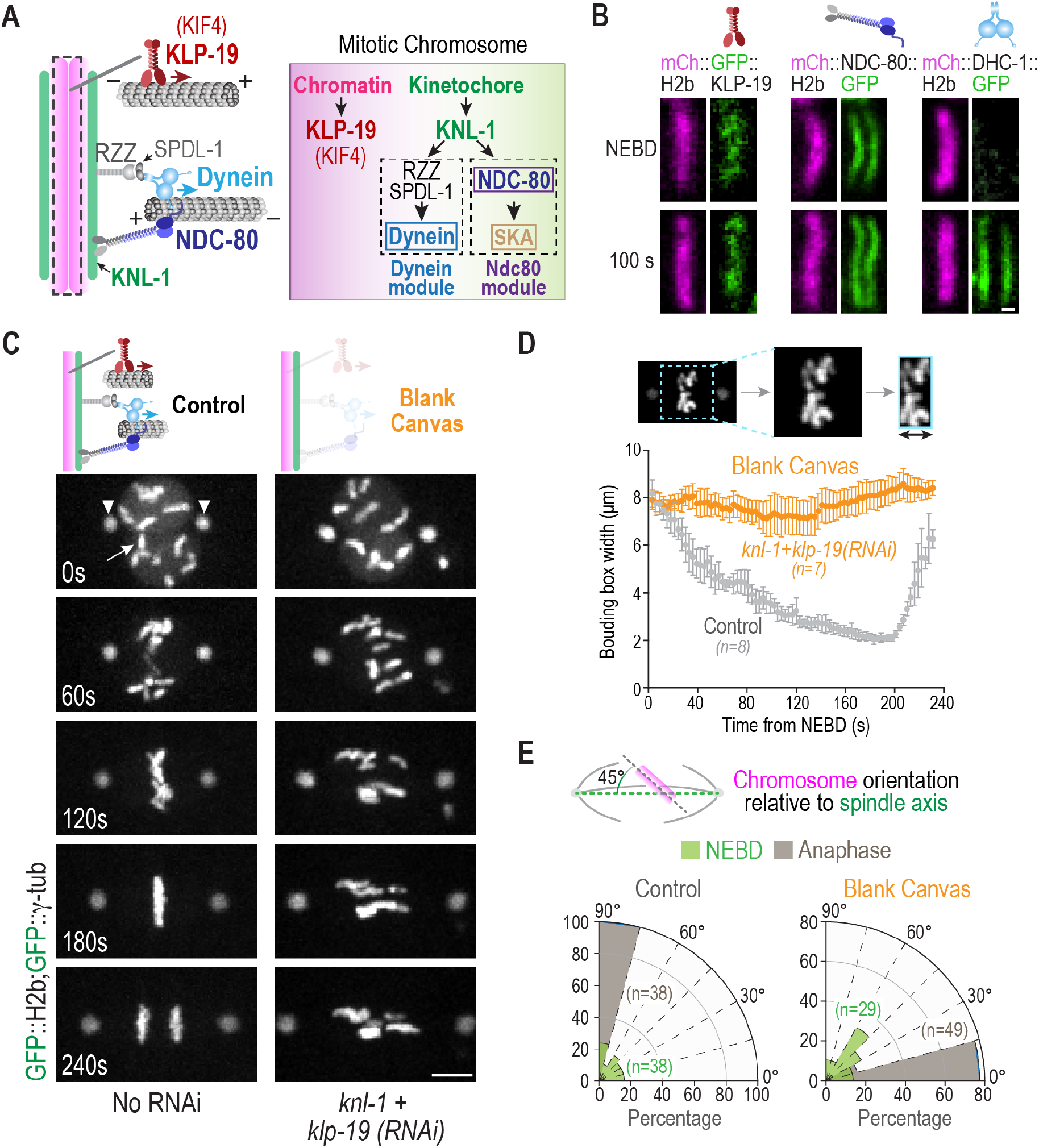
Creating a blank canvas on mitotic chromosomes with respect to spindle microtubule interactions in the *C. elegans* embryo. **(A)** Schematics of the three major microtubule-targeting factors on mitotic chromosomes (*left*) and their assembly dependencies (*right*) in the one-cell *C. elegans* embryo. **(B)** Chromosomal localization of *in situ*-tagged GFP fusions of the indicated components. Scale bar, 0.5 µm. **(C)** Phenotype of the blank canvas state generated by removing the chromokinesin KLP-19 and preventing outer kinetochore assembly by depletion of KNL-1. GFP fusions of histone H2b and γ-tubulin label the chromosomes (*arrow*) and spindle poles (*arrowheads*), respectively. Scale bar, 5 µm. **(D)** Minimal bounding box analysis quantifying chromosome dispersion on the spindle. Graph plots the mean bounding box width following NEBD for the indicated conditions. Error bars are the 95% confidence interval. **(E)** Quantification of chromosome orientation relative to the spindle pole-to-pole axis at NEBD and anaphase onset. Radial plots show the percentage of chromosomes within 15° angular orientation bins. Anaphase onset in the blank canvas state, where chromosome segregation fails, was scored by initiation of spindle rocking. *n* represents the number of chromosomes measured per condition.

Cytoplasmic dynein and the Ndc80 module are both recruited to kinetochores (**Fig. 1A,B**), the former via the Rod/Zwilch/Zw10 (RZZ) complex and the activating adapter Spindly (SPDL-1 in *C. elegans*), and the latter via the kinetochore linker & scaffold Mis12 and Knl1 complexes ^3, 12^; in many species, including vertebrates, the Ndc80 module is additionally recruited by CENP-T ^13–15^. While the Ndc80 complex assembles at kinetochores prior to nuclear envelope breakdown (NEBD), dynein is recruited only after NEBD (**Fig. 1B**); in *C. elegans*, the Ska complex is recruited significantly later and its recruitment requires Ndc80 complex engagement with the microtubule lattice ^5^. In contrast to these kinetochore-localized microtubule-targeted activities, chromokinesins are broadly recruited to mitotic chromatin, potentially via direct binding to DNA and via interaction with condensin complexes ^1, 16–20^ (**Fig. 1A,B**).

While significant phenotypic and biochemical/structural analysis has been conducted for these conserved chromosome segregation factors in different species, the complexity of their coordinated action has limited understanding of their individual contributions in an *in vivo* context. We therefore decided to pursue a *in vivo* reconstruction approach, in which we first characterized the effect of removing all three major force generators, creating a blank canvas on mitotic chromosomes with respect to microtubule interactions. We then compared chromosome dynamics in the blank canvas state to specifically engineered conditions where individual force generators were selectively present on mitotic chromosomes. We conducted this analysis in the early *C. elegans* embryo, where conserved chromosome segregation factors have been extensively investigated, the kinetochore assembly hierarchy *in vivo* is well-characterized, kinetochore composition is relatively streamlined, and the diffuse line-shaped kinetochore geometry enables facile and dynamic readout of chromosome orientation on the spindle.

To generate a blank canvas on mitotic chromosomes, we co-depleted KNL-1, which is essential for outer kinetochore assembly in *C. elegans*, and the KIF4 family chromokinesin KLP-19, which is the major mitotic chromatin-localized motor in this system ^17, 21^. In wild-type embryos, the 12 chromosomes from the two pronuclei are rapidly aligned and congressed, with anaphase onset occurring ∼3 min after NEBD. By contrast, in the blank canvas state, after NEBD the chromosomes remained dispersed on the spindle and became aligned parallel to the spindle pole-to-pole axis (**Fig. 1C**). To quantify chromosome dynamics, we employed two measures: a bounding box that measures the width of the dispersion of all chromosomes on the spindle ^5^ (**Fig. 1D**), and the angular orientation of chromosomes relative to the spindle pole-to-pole axis at the time of NEBD and anaphase onset (**Fig. 1E**). The bounding box analysis indicated that in the blank canvas state there was no significant congression of chromosomes towards the spindle equator (**Fig. 1D**). The angular orientation analysis indicated that, in contrast to the perpendicular orientation at the time of anaphase onset in controls, chromosomes exhibited parallel orientation relative to the spindle pole-to-pole axis in the blank canvas state (**Fig. 1E**). Thus, in the absence of chromatin and kinetochore-localized force generators/couplers, mitotic chromosomes are distributed throughout the spindle and have their axes oriented parallel to the spindle pole-to-pole axis. The parallel orientation of chromosomes likely arises through nematic alignment caused by the action of dynamic microtubule polymers pushing on chromosomes as passive objects.

The generation and characterization of a blank canvas with respect to mitotic chromosome-microtubule interactions prompted us to next engineer *in vivo* states in which only one of the major force generators is present on mitotic chromosomes (**Fig. 2A**). To create a chromokinesin-only state, we depleted KNL-1 (“ChrKin only”); to create a Ndc80 module-only state, we co-depleted KLP-19 and ROD-1, a subunit of the RZZ complex that recruits dynein to kinetochores (“Ndc80 only”); to create a kinetochore dynein-only state, we co-depleted KLP-19 and NDC-80 in a strain harboring an RNAi-resistant transgene expressing a mutant form of NDC-80 that disrupts the ability of its conserved calponin homology (CH) domain to dock onto the microtubule surface but does not otherwise affect the complex ^5, 22^ (“Dynein only”) (**Fig. 2A**). The use of this mutant form of NDC-80, which has been rigorously characterized in prior work^5, 23^, minimizes impact on kinetochore structure. In all cases, we monitored chromosome distribution on the spindle and axial orientation of chromosomes relative to the spindle axis, and compared the outcomes to the blank canvas, where none of these force generators are present on chromosomes.

**Figure 2.**
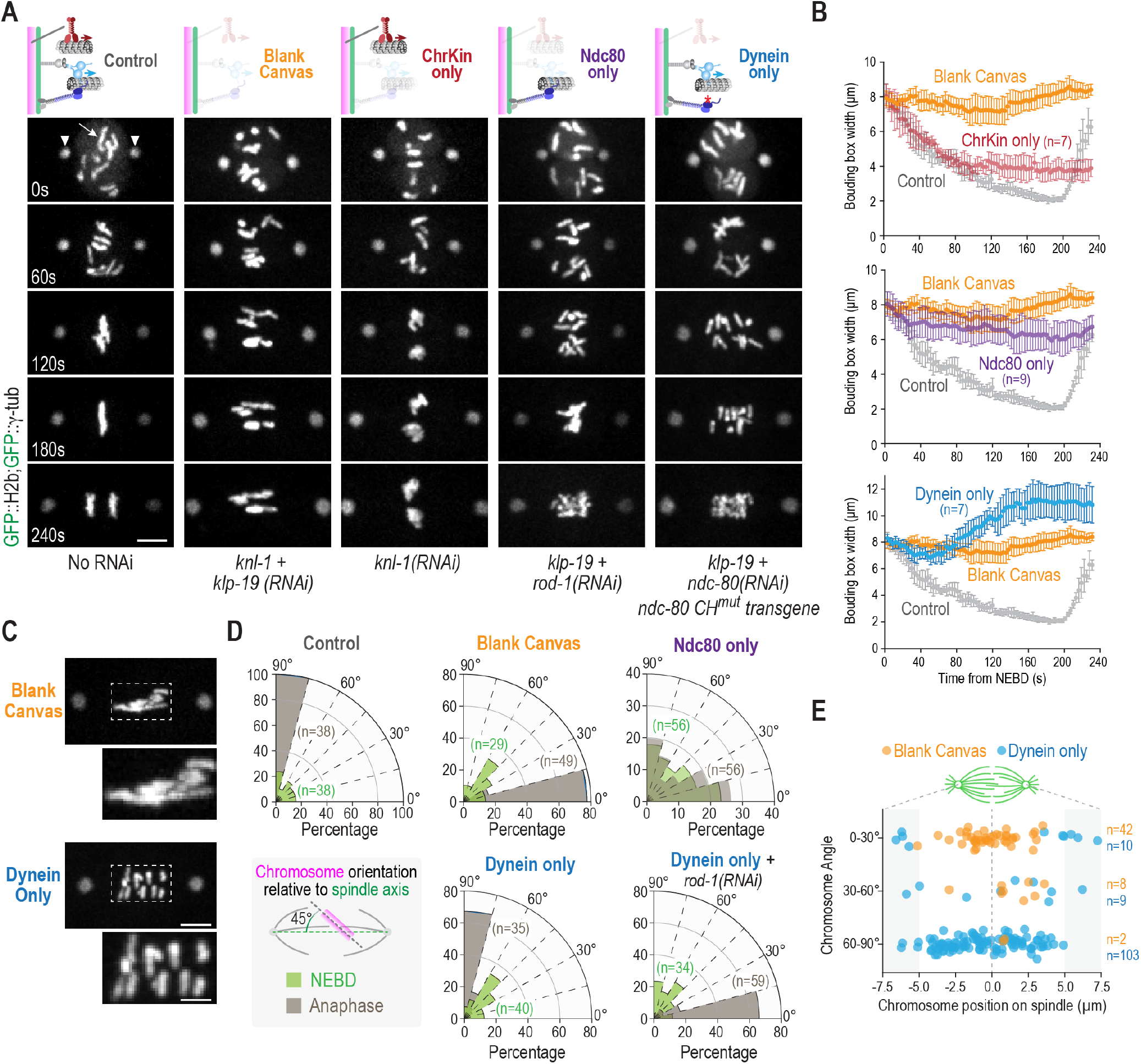
An *in vivo* reconstruction approach to define the contributions of individual microtubule-directed activities on mitotic chromosomes. **(A)** Image panels from timelapse movies monitoring chromosomes (*arrow*) and spindle poles (*arrowheads*) for the indicated conditions. Time relative to NEBD is indicated in the lower left of the Control panels. Schematics (*top*) indicate the specific states created and analyzed; label below each panel column indicates the perturbation(s) used to create that state. Scale bar, 5 µm. **(B)** Quantification of chromosome dispersion on the spindle performed as in Fig. 1D. The Control and Blank Canvas curves are the same as in Fig. 1D and are plotted with specific perturbations to aid comparison. *n* is the number of embryos analyzed. Error bars are the 95% confidence interval. **(C)** Representative images of the blank canvas and kinetochore dynein only states, highlighting the striking difference in chromosome orientation. Scale bar for spindle panels, 5 µm; boxed region is magnified 2X below, scale bar 2.5 µm. **(D)** Quantification of chromosome orientation relative to the spindle axis, similar to Fig. 1E. *n* is the number of chromosomes analyzed per condition. **(E)** Plot of chromosome angle relative to chromosome position on the spindle, comparing the blank canvas and kinetochore dynein only states. The spindle equator is the origin for the x-axis and is indicated with a dashed line. *n* is the number of chromosomes measured.

In the presence of only the chromokinesin KLP-19, which is equivalent to the previously described “kinetochore null” phenotype ^21^, the chromosomes from the oocyte and sperm nuclei moved to the spindle center, formed tight balls lacking any discernable orientation, and no segregation was observed (**Fig. 2A**). Thus, chromokinesin activity is sufficient to move chromosomes towards the spindle equator (**Fig. 2B**), but the chromosomes from each pronucleus are tightly compressed together with no apparent orientation or segregation; they also do not achieve the same degree of tight congression observed in wildtype embryos. In the presence of the Ndc80 module only, where the Ndc80 and Ska complexes are present but kinetochore dynein and chromokinesin are not, chromosomes exhibit delayed partial congression and extensive mis-orientation, leading to bridging during anaphase segregation (**Fig. 2A,B,D**). Analysis of chromosome distribution highlighted the late partial congression supported by the Ndc80 module (**Fig. 2B**), which in the wildtype may reflect engagement of this module after primarily chromokinesin-driven chromosome movement towards the spindle equator.

The most striking chromosome behavior observed using the *in vivo* reconstruction approach was in the kinetochore dynein-only state. Like the blank canvas, chromosomes remained dispersed on the spindle in the kinetochore dynein-only state (**Fig. 2A,B**), indicating that kinetochore dynein is unable to support significant congression. However, in stark contrast to the blank canvas state, where chromosomes were uniformly aligned parallel to the spindle axis, the majority of chromosomes were oriented perpendicular to the spindle axis by the time of anaphase onset (**Fig. 2A,C,D**). Plotting the angular orientation of chromosomes relative to their position on the spindle indicated the presence of a small number of pole-proximal chromosomes that failed to orient in the dynein-only state and remained aligned parallel to the spindle axis (**Fig. 2E**); the most likely explanation for the orientation of these chromosomes is that minus end-directed kinetochore dynein motor activity, in the absence of chromokinesin, traps them in the high density parallel-oriented microtubule environment close to the spindle poles. The presence of these pole-trapped chromosomes accounts for the greater average chromosome dispersion on the spindle in the kinetochore dynein-only state relative to the blank canvas state (**Fig. 2B**). To confirm that the striking change in chromosome angular orientation was indeed due to the action of kinetochore dynein, we depleted ROD-1 in the condition employed to generate the kinetochore dynein-only state. Co-depletion of ROD-1 resulted in chromosomes once again exhibiting the parallel alignment observed in the blank canvas state (**Fig. 2C,D; Fig. S1A**), indicating that RZZ complex-dependent dynein recruitment to kinetochores drives the change in chromosome orientation. In support of this conclusion, removal of dynein from microtubule plus ends by deletion of the plus end-binding protein EBP-2 ^24^ did not affect chromosome orientation in the kinetochore dynein-only state (**Fig. S1B**). Thus, kinetochore dynein recruited in a RZZ complex-dependent manner is sufficient to drive the striking transition in chromosome angular orientation relative to the spindle axis. A comparison of the conditions where only the Ndc80 module or the kinetochore dynein module is present is informative (**Fig. 2A-D**). The Ndc80 module drives late, partial congression but is unable to ensure proper chromosome orientation. By contrast, the dynein module is unable to drive any congression but is remarkably efficient at orienting chromosomes.

The striking change in chromosome orientation observed in the kinetochore dynein-only state, despite their dispersion on the spindle, suggested that kinetochore dynein was sufficient not only for chromosome angular orientation relative to the spindle axis but also for biorientation, the state in which sister chromatids attach to opposite spindle poles. To assess if this was indeed the case, we imaged *in situ*-tagged KNL-1::GFP, which marks the diffuse sister kinetochores on individual chromosomes as paired lines. Imaging of KNL-1::GFP revealed that, in the kinetochore dynein-only state, the majority of sister kinetochore pairs (92%) were indeed bioriented and faced opposite spindle poles (**Fig. 3A**). The biorientation was achieved by a combination of rotation and translational motions of individual chromosomes (**Fig. 3B**). In addition, despite their dispersion throughout the spindle, sister chromatids exhibited aspects of separation towards opposite spindle poles upon anaphase onset (**Fig. 3C**). While not as uniform and persistent as in the wildtype, this separation is consistent with biorientation of sister kinetochores. We note that removal of the kinetochore dynein module has been associated with chromosome missegregation in *Drosophila*, *C. elegans* and human cells ^25–27^, with the *Drosophila* observations being over three decades old. However, the reasons for this missegregation have remained unclear. The results above suggest that loss of a direct contribution to chromosome orientation is a contributing factor to the missegregation and developmental lethality caused by loss of kinetochore dynein. At least in *C. elegans* and *Drosophila*, we can exclude contribution to lethality and missegregation from perturbation of the spindle checkpoint caused by RZZ complex inhibition ^28–30^ (**Fig. S2A,D,E**). The rapid timescales involved in the *C. elegans* embryo also make it unclear to what extent compromising fibrous corona expansion that is observed on persistently unattached kinetochores and requires the RZZ complex ^31^, is a contributing factor. We additionally note that the concerted action of the Ndc80 module and chromokinesin, in the absence of kinetochore dynein, is able to congress and biorient a substantial proportion of chromosomes (**Fig. S2B,C**) – thus, with respect to the biorientation function of chromosomal microtubule-targeted activities, the chromokinesin and Ndc80 module combination can be considered as acting in parallel to the kinetochore dynein module, although action of all three is required to ensure that every chromosome is properly bioriented and accurately segregated. The comparative analysis additionally indicates that the chromokinesin and Ndc80 module combination drives chromosome congression, with no significant contribution from kinetochore dynein (**Fig. S2C**). Parallel action of kinetochore dynein and the chromokinesin-Ndc80 module combination likely explains why observing the sufficiency of kinetochore dynein for chromosome biorientation required establishment of the *in vivo* reconstruction approach.

**Figure 3.**
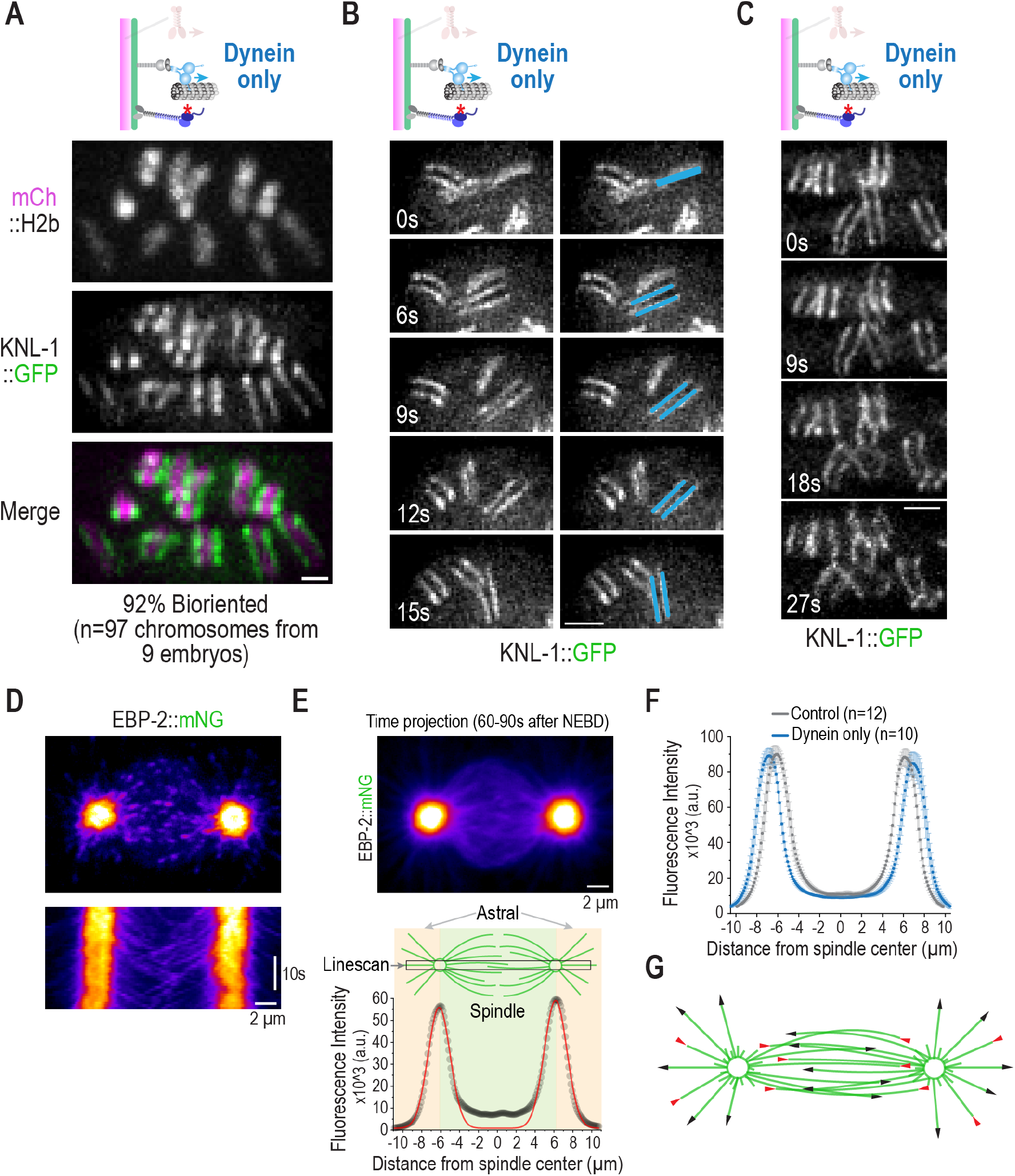
Biorientation in the kinetochore dynein only state and spindle microtubule behavior during the time of chromosome orientation. **(A)** Representative single channel and 2-color overlay images of a kinetochore marker and fluorescent histone H2b. Quantification of biorientation is indicated below the image panels. Scale bar, 1 µm. **(B)** (*left*) Dynamics of a chromosome flip visualized using the kinetochore marker KNL-1::GFP. The first frame was arbitrarily set to 0s. (*right*) The same image panels with one specific orienting chromosome highlighted. Scale bar, 2 µm. **(C)** Visualization of anaphase-like sister chromatid movements, despite lack of congression, in the dynein only state. Scale bar, 2 µm. **(D)** Rapid (every 0.8s) single plane imaging of EBP-2 dynamics starting ∼1 min after NEBD. Image on top shows a single frame highlighting EBP-2 comets at microtubule plus ends and kymograph below shows rapid loss of EBP-2 signal moving out from the spindle pole center and tracks of EBP-2 comets invading opposing half-spindles. Scale bar, 2 µm for both. **(E)** (*top*) Time projection (30.4 seconds/38 frames) of a representative EBP-2 sequence starting ∼1 min after NEBD; Scale bar, 2 µm. (*bottom*) Linescan analysis of the sum projection image using a 1.5 x 22 µm box drawn as indicated in the spindle schematic. The intensity profile of the linescan was fit to a double Gaussian (*red line*). **(F)** Analysis of EBP-2 dynamics as in *(E)*, for both control and dynein only states. *n* is the number of embryos imaged. Error bars are the 95% confidence interval. **(G)** Schematic summarizing spindle microtubule behavior extrapolated from EBP-2 analysis in the ∼1-1.5 min interval after NEBD, when chromosome orientation is observed.

The cytoplasmic dynein motor complex, together with its motility co-factor dynactin and activating adapters such as Spindly, exhibits minus end-directed motility along the microtubule lattice^32^. Such motility would lead to chromosomes moving poleward with kinetochore dynein laterally bound to the microtubule surface and has been observed on rare occasions during initial capture of spindle microtubules by kinetochores in vertebrate cells ^33, 34^. However, minus end-directed lattice motility is unlikely to explain the striking biorientation of the majority of chromosomes driven by the kinetochore dynein module. Biophysical analysis has shown that when a depolymerizing microtubule end reaches a lattice-bound dynein molecule, the depolymerization of the microtubule is suppressed and the force from the depolymerizing end transmitted to the object coupled to the motor ^35, 36^. Given the outward peeling geometry of protofilaments of depolymerizing microtubules, such an end-coupled dynein interaction would exert a torque on the bound object ^37^, such as a chromosome. However, for the chromosome to rotate and orient towards the pole, there has to be a significant likelihood for converting lattice-bound motile states to end-coupled dynein attachments. To address the feasibility for achieving an end-coupled state of already-laterally-associated kinetochore dynein, we analyzed microtubule behavior in the spindle using the plus end-tracking protein EBP-2 ^38^ (**Fig. 3D**), in both controls and the kinetochore dynein-only state. While metaphase spindle structure, in the one cell *C. elegans* embryo has been analyzed in-depth by 3D EM reconstruction ^39^, our analysis focused earlier (∼1 min after NEBD) when chromosome-microtubule attachments are being established and chromosomes begin biorienting and congressing. Kymographs and time projections of fast-acquisition single plane EBP-2 image series revealed rapid decay of EBP-2 signal close to the spindle poles (**Fig. 3D,E**). However, once past an ∼2 µm zone around the pole, microtubules on the spindle side extended significantly past the spindle equator and into the opposing half spindle (**Fig. 3D-F**); such EBP-2 behavior is consistent with the notion that a nuclear factor, such as Ran-GTP, biases microtubule growth on the spindle side ^40, 41^. As spindle poles are known to concentrate microtubule depolymerizing activity ^42, 43^, these observations suggest a picture in which microtubules on the spindle side that escape the ∼2 µm polar zone grow persistently until reaching the vicinity of the opposite spindle pole (**Fig. 3G**). The penetration of microtubules from one pole into the opposing half spindle may help ensure that initial lateral capture of microtubules by kinetochore dynein at different spindle positions will convert with some probability into an end-coupled state (this probability depends on the lifetime of the kinetochore dynein-lattice interaction versus the lifetime of the bound microtubule). We speculate that a transition from laterally associated to an end-bound dynein state on an individual kinetochore is central to the biorientation function of kinetochore dynein. As chromosomes frequently translocate poleward as they orient (see *Fig. 4B,C* below), the proposed kinetochore dynein end-bound state may also couple to depolymerization or cooperate with other kinetochore dynein modules on the same kinetochore engaged in minus end motility. The extremely fast dynamics (orientation occurs in ∼10-15s time frame) and the absence of discrete kinetochore fibers makes dynamically visualizing the geometry of kinetochore-microtubule interactions in *C. elegans* embryos currently unfeasible. One expectation from this line of thinking is that orientation will be kinetochore-autonomous, with a single kinetochore orienting the chromosome independently of its sister. To assess if this is the case and to gain insight into the role of kinetochore dynein in sensing microtubule attachments at kinetochores, we next focused on coordinated analysis of chromosome orientation and outer kinetochore remodeling.

**Figure 4.**
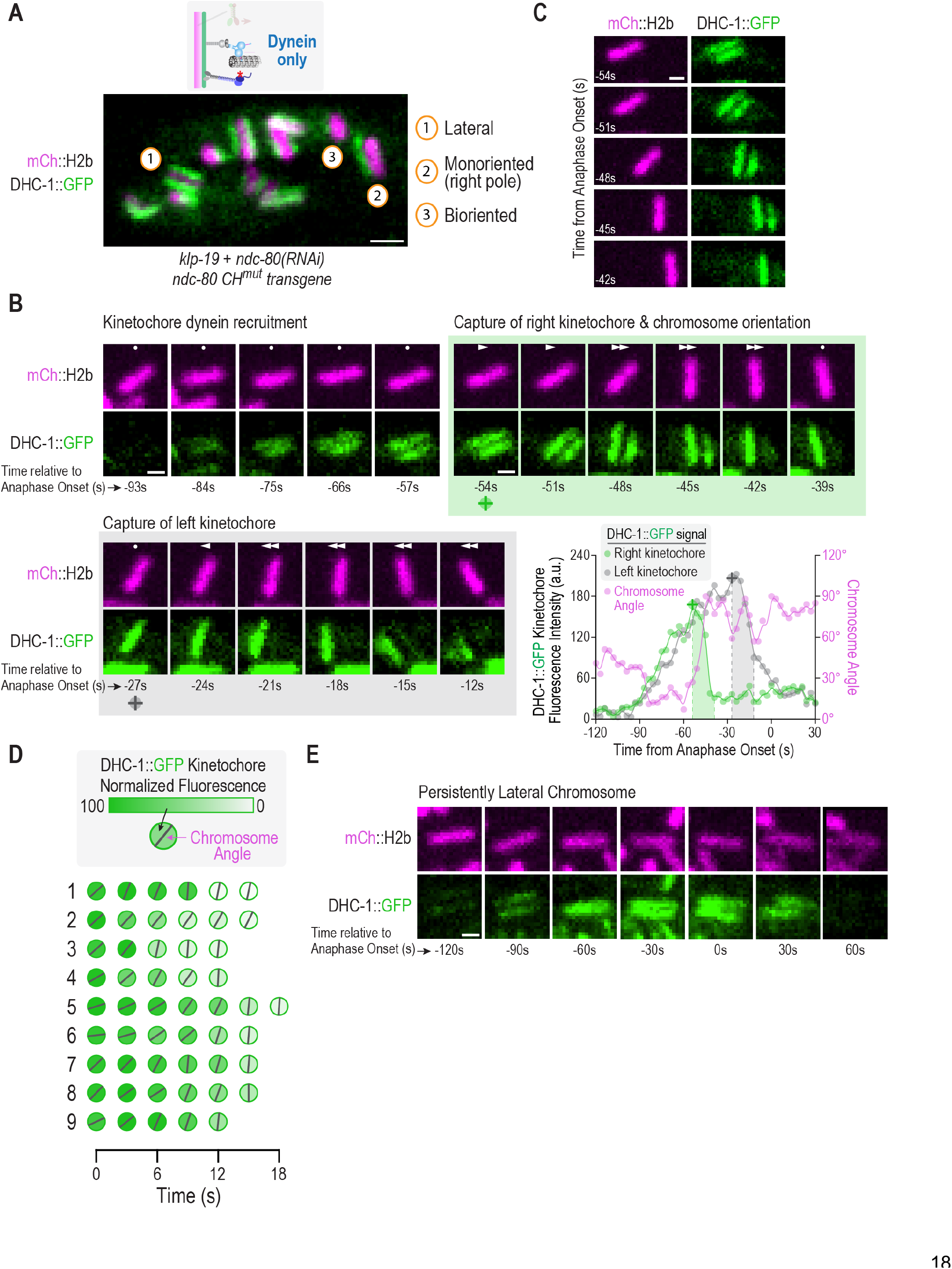
Orientation-coupled removal of dynein from kinetochores in the dynein only state. **(A)** Snapshot of dynein localization, visualized using *in situ*-tagged DHC-1::GFP, at a time point when chromosomes are autonomously orienting on the spindle in the kinetochore dynein only state. Numbered circles highlight 3 chromosomes in different states. **(B)** Image sequence of a single chromosome and of dynein localized to its two sister kinetochores. To aid comparison, the panels are centered on the chromosome; dots (*no movement*) and arrows (*directional movement*) in each panel indicate chromosome movement relative to the spindle pole (*see also panel C*). Green and grey shaded boxes highlight successive capture of the two sister kinetochores. Times are relative to anaphase onset in seconds. Graph on the lower right quantifies the DHC-1::GFP fluorescence intensity on each kinetochore, along with the chromosome angle relative to the pole-pole axis. Shaded areas on the graph correspond to the shaded boxes around image panel sets. Symbols below −54s and −27s panels serve as reference points linking image panels to the graph. Scale bar, 1 µm. **(C)** Same chromosome as in *(B)* during the time of right kinetochore capture and orientation, highlighting poleward translocation of the chromosome during this process. Scale bar, 1 µm. **(D)** Analysis of kinetochore dynein fluorescence intensity and chromosome angle for 9 kinetochores. Shading in the circle indicates DHC-1::GFP fluorescence signal and the black line indicates the chromosome angle relative to the spindle axis. For simplicity, translocation on the spindle is not shown. **(E)** DHC-1::GFP localization on a chromosome that maintains a persistent lateral orientation through anaphase and does not biorient. Scale bar, 1 µm.

A well-established function of the kinetochore dynein module is to remodel the outer kinetochore following the establishment of end-coupled attachments. Specifically, minus end motility of kinetochore dynein removes the spindle checkpoint-activating Mad1-Mad2 complexes following establishment of end-on attachments in order to silence checkpoint signal generation and promote cell cycle progression ^44, 45^. Notably, a majority of kinetochore dynein is itself removed along with checkpoint components, with puncta of kinetochore dynein module components and associated checkpoint proteins observed moving towards spindle poles ^44, 46^. The precise mechanisms by which the removal event is triggered is unclear, aside from a requirement for end-on attachment; removal is not observed when kinetochores are laterally attached to microtubules ^47^. One potential model is that Ndc80 complex engagement with microtubule ends is a pre-requisite to trigger the removal (Ndc80 sensor model). A second model is that end-coupled interactions deliver a regulatory activity, such as a protein phosphatase, that triggers dissociation between elements of the kinetochore dynein module and/or between the module and its binding interface on the outer kinetochore (Phosphatase delivery model).

To distinguish between these and other models, and to gain more insight into the biorientation function of kinetochore dynein, we imaged *in situ* GFP-tagged dynein heavy chain (DHC-1::GFP), in the kinetochore dynein-only state. A snapshot ∼100s after NEBD revealed a heterogeneous population of chromosomes, all with different positions and orientations (**Fig. 4A**), consistent with orientation being a chromosome-autonomous process. The amount of DHC-1 on kinetochores varied widely between individual chromosomes and correlated with chromosome orientation. Both sister kinetochores of laterally oriented chromosomes were enriched for dynein, but perpendicularly oriented chromosomes had dynein concentrated on one or neither of their sister kinetochores (**Fig. 4A**).

To capture the detailed temporal relationship between chromosome orientation and dynein dynamics at the kinetochore, we imaged chromosomes and DHC-1 at high temporal resolution (**Fig. 4B**). A single chromosome that exhibited all of the distinct phases of DHC-1 dynamics and achieved biorientation is shown in **Fig. 4B**. Following NEBD, both sister kinetochores on this chromosome lacked DHC-1 but then began to recruit it simultaneously; the chromosome was oriented parallel to the spindle pole-to-pole axis at this time. Soon after, the chromosome rotated while translocating towards the right pole, which coincided with abrupt loss of dynein from the kinetochore facing that pole (**Fig. 4B,C**). During the orientation, poleward translocation, and dynein removal from the right kinetochore, there was no reduction in dynein levels on the left sister kinetochore (**Fig. 4B,C**), highlighting that orientation and dynein removal are kinetochore-autonomous events that primarily reflect the engagement of one kinetochore with one spindle pole. Next, the left sister kinetochore was captured by the left pole, causing leftward translocation, and leading to eventual loss of a large proportion of dynein from this kinetochore (**Fig. 4B**). Quantification of the amount of dynein and chromosome angle relative to the spindle pole-to-pole axis for this chromosome reveals; 1) the similar and continuous buildup of dynein on both kinetochores during the initial phase, 2) the abrupt loss of the majority of dynein on the right sister kinetochore coincident with orientation of the chromosome perpendicular to the spindle axis, and 3) subsequent capture and loss of dynein from the left sister kinetochore, and 4) a residual pool of dynein that likely supports kinetochore-microtubule interactions in the bioriented state (**Fig. 4B**). Data for 9 kinetochores monitored similarly in the kinetochore dynein-only state are shown in **Fig. 4D** and they all indicate coupling of orientation and dynein removal. In contrast to chromosomes that achieved the perpendicular orientation, rare chromosomes that were trapped in a lateral orientation (**Fig. 2E**) maintained consistent dynein levels at both sister kinetochores until anaphase onset (**Fig. 4E**). Kinetochore dynein dissociated from these chromosomes soon after anaphase onset, suggesting that its loss is triggered by a global change in cell cycle state rather than a specific microtubule attachment configuration. This contrast between laterally oriented versus perpendicularly bioriented chromosomes lends additional support to the proposal that an end-coupled dynein attachment state underlies both the orientation and the removal. In addition to DHC-1, we also imaged the RZZ complex subunit ROD-1 and the kinetochore dynein activator SPDL-1. SPDL-1 behaved similarly to DHC-1 whereas ROD-1 was not removed coincident with kinetochore-autonomous orientation (**Fig. S3A,B**). These observations suggest the removal event primarily involves dissociation between the RZZ complex and the rest of the kinetochore dynein module, along with its cargo. In addition, the fact that maintaining biorientation in the dynein-only state requires ROD-1 (**Fig. 2D, Fig. S1A**), suggests that a small stable pool of RZZ-associated dynein persists after biorientation. The most prominent cargo for removal by the kinetochore dynein module is the spindle checkpoint-activating Mad1-Mad2 complex. We therefore also imaged in situ-tagged MAD-1 in the kinetochore dynein-only state. Unfortunately, high signal for MAD-1 in the general spindle area along with its significantly slower recruitment relative to DHC-1 made imaging and quantifying its kinetochore dynamics in the dynein-only state challenging (robust analysis of MAD-1 localization in *C. elegans* embryos requires generation of monopolar spindles ^48^). However, qualitatively MAD-1 behavior was similar to that of DHC-1 (**Fig. S3C**). Overall, the results indicate that chromosome orientation and outer kinetochore remodeling are tightly coupled in the kinetochore dynein-only state.

As the NDC-80 complex’s microtubule-binding activity is inactivated through mutation in the kinetochore dynein-only state, the above data argue against an Ndc80 sensor model in which microtubule lattice engagement of Ndc80 is the cue for initiating the removal reaction. In support of this conclusion, comparison of the NDC-80 microtubule-binding mutant to NDC-80 WT, effectively comparing kinetochore dynein-only to kinetochore dynein plus NDC-80 module states, revealed no significant difference in DHC-1 removal from kinetochores following chromosome orientation (**Fig. 5A**). In this analysis, we measured DHC-1 signal at both sister kinetochores before and after one sister oriented towards one pole. In addition to indicating robust removal of DHC-1 regardless of Ndc80 status, the analysis further confirmed the kinetochore-autonomous removal of DHC-1 (**Fig. 5A**); the interval between the before and after measurement timepoints, which reflects the rate of removal, was also unaffected (**Fig. S3D**). This conclusion is consistent with RNAi analysis of the Ndc80 complex in human cells, where significant Ndc80 depletion does not prevent removal of spindle checkpoint components^49^.

**Figure 5.**
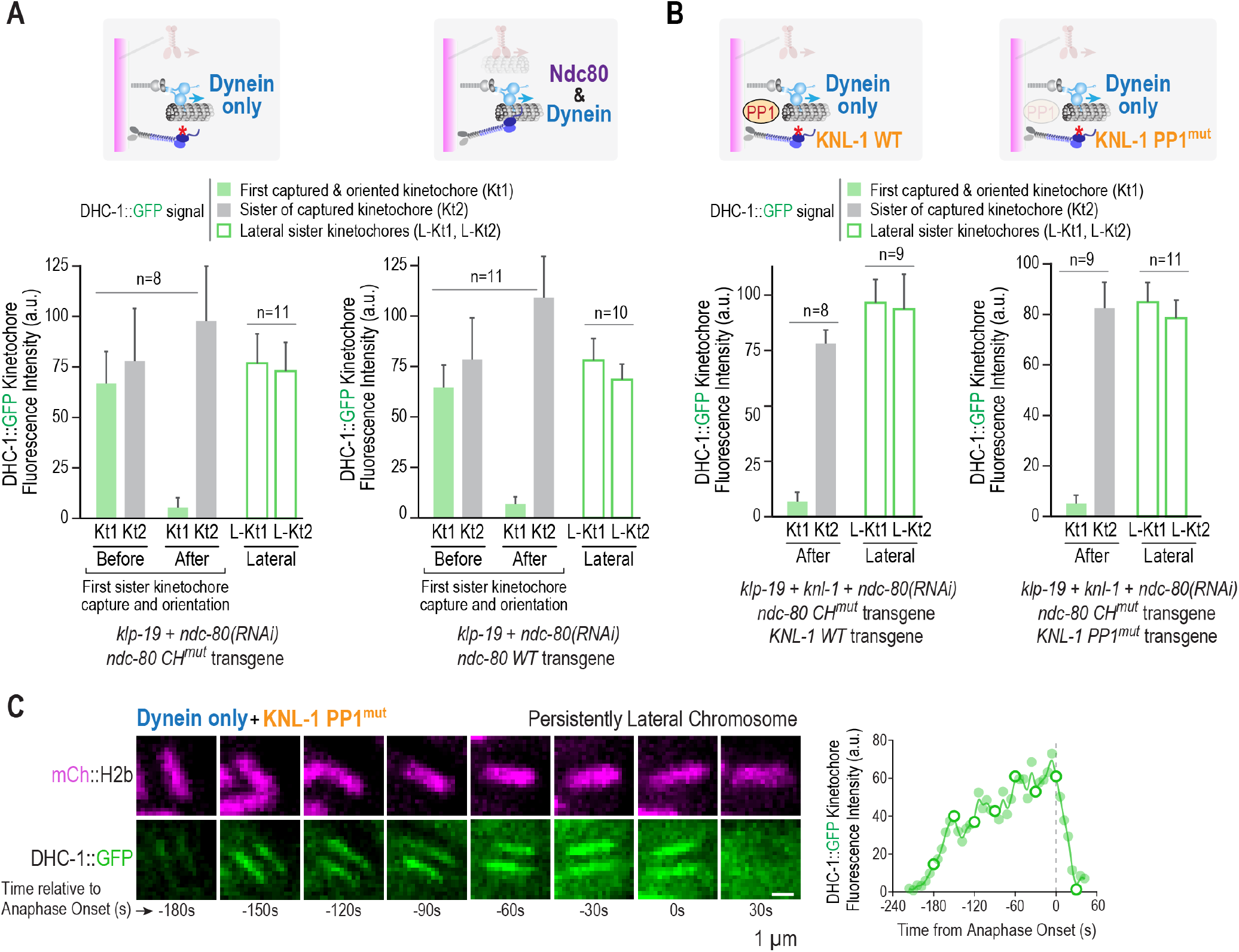
Dynein removal from kinetochores is independent of the Ndc80 module and kinetochore-localized protein phosphatase 1. **(A)** Comparison of kinetochore dynein removal with a functional or mutant Ndc80 module. Schematics on top indicated the states analyzed; text below the graphs indicates the perturbations used to generate these states. DHC-1::GFP was quantified on sister kinetochores before and after one sister was captured and oriented towards a pole; Kt1 and Kt2 refer to the captured kinetochore and its sister, respectively. DHC-1::GFP on sister kinetochores of chromosomes that maintained a persistently lateral orientation until anaphase onset was also measured (L-Kt1, L-Kt2). *n* is the number of sister kinetochore pairs (chromosomes) analyzed. **(B)** Analysis of kinetochore dynein removal with or without kinetochore-localized protein phosphatase 1 (PP1). Schematics on top indicated the states analyzed; text below the graphs indicates the perturbations used to generate these states. Analysis was as in *(A)*, except that DHC-1::GFP signal was measured on sister kinetochores after but not before chromosome orientation. **(C)** Loss of DHC-1::GFP from kinetochores of persistently lateral chromosomes following anaphase onset. Graph on right plots the average signal for both kinetochores of the imaged chromosome. Open circles on graph correspond to the time points shown in the image panels. Scale bar, 1 µm.

We next tested if protein phosphatase 1 (PP1) delivery to kinetochores is important for triggering DHC-1 removal. PP1 localization at kinetochores is complementary to that of checkpoint proteins ^50^, suggesting that it could serve as a removal trigger. For this purpose, we combined the kinetochore dynein-only state with manipulation of KNL-1, the primary PP1-targeting protein at kinetochores in the *C. elegans* embryo ^51, 52^. We compared WT KNL-1 to a well-characterized PP1 docking mutant ^51, 52^ (PP1^mut^) but did not observe any significant difference between these two conditions (**Fig. 5B; Fig. S3D**). The PP1 docking mutant also did not affect removal of DHC-1 from the kinetochores of rare persistently lateral chromosomes after anaphase onset (**Fig. 5C; Fig. S3E**). These results argue against a PP1 trigger model but leave open the possibility that another regulatory activity initiates removal of the kinetochore dynein module and associated checkpoint activators; however, any such activity must be sensitive to orientation/attachment state. Alternatively, removal may be intrinsic to the kinetochore dynein module once it achieves a specific attachment state, most likely an end-coupled state. We favor the latter possibility given the ability of the dynein module to orient chromosomes and the tight coupling between orientation and removal. Overall, the results argue against Ndc80 sensor and PP1 delivery models for triggering removal and highlight tight coupling between kinetochore-autonomous orientation and remodeling by the kinetochore dynein module.

In summary, the establishment of an *in vivo* reconstruction approach, in which the presence of individual microtubule-targeting activities on mitotic chromosomes is compared to a blank canvas state, revealed that the kinetochore dynein module is sufficient to biorient chromosomes on the spindle and support remodeling of the outer kinetochore; both of these actions occur in a chromosome-autonomous and kinetochore-autonomous manner. This activity of the kinetochore dynein module acts largely in parallel to the combined action of chromokinesin and the Ndc80 module, accounting for the moderate orientation and segregation defects observed when the dynein module is absent. Notably, the segregation defects associated with loss of kinetochore dynein lead to lethality, indicating that even when the other activities are present, the function of the kinetochore dynein module is essential for viability. While dynein in the context of the fibrous corona of the kinetochore has long been proposed to aid lateral capture of microtubules ^34^, the findings here, along with prior work ^23^, suggest that an end-coupled state of dynein is functionally significant for both orientation and remodeling activities of this conserved motor module at metazoan kinetochores. To date, only two studies have characterized the biophysical properties of the end-coupled dynein state *in vitro* ^35, 36^, in contrast to the myriad biophysical studies of dynein motility. Directly visualizing and manipulating the proposed end-coupled dynein state, which we suggest plays a critical role during mitotic chromosome segregation and may also contribute to other dynein functions, will be an important future goal.

## ACKNOWLEDGMENTS

We thank Reto Gassmann and François Nédélec for helpful discussions and Sander van den Heuvel for sharing strains. This work is supported by an NIH grant to A.D. (R01 GM074215), a Sir Henry Wellcome Postdoctoral Fellowship (215925) to B.P. and a Sir Henry Dale Fellowship (208833) to D.K.C. Currently, B.P. is sponsored by W.C.E. A.D. acknowledges salary support from the Ludwig Institute for Cancer Research.

## AUTHOR CONTRIBUTIONS

B.P. and A.D. conceived the study. B.P., D.K.C. and A.D. designed the experiments. B.P. conducted and analyzed the experiments. B.P. and D.K.C. generated the *C. elegans* strains. B.P. and A.D. prepared the manuscript.

## SUPPLEMENTAL FIGURES & LEGENDS

**Figure S1.**
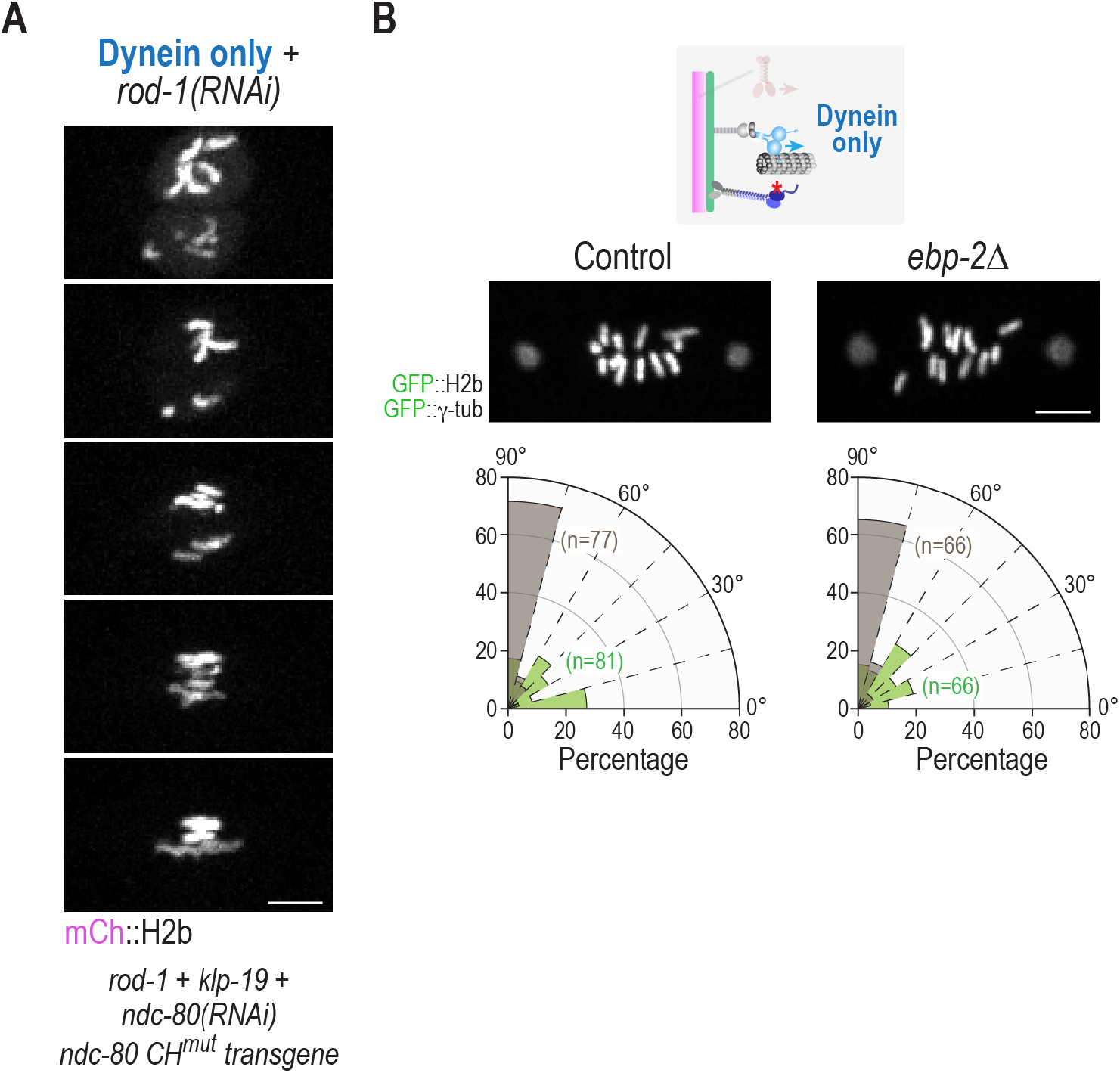
Representative image sequence of ROD-1 depletion in the dynein only state and analysis of *ebp-2Δ*. **(A)** Representative images from a timelapse series following removal of ROD-1 in the condition used to generate the dynein only state. Text below the panel indicates the specific perturbations used in this experiment. Quantification of chromosome angle in this condition is shown in Fig. 2D. Scale bar, 5 µm. **(B)** Analysis of the kinetochore dynein only state in control or *ebp-2Δ* embryos. Loss of EBP-2, which prevent plus end tracking of dynein, has no significant effect on chromosome orientation, indicating that RZZ-recruited dynein is responsible for the orientation function. Scale bar, 5 µm.

**Figure S2.**
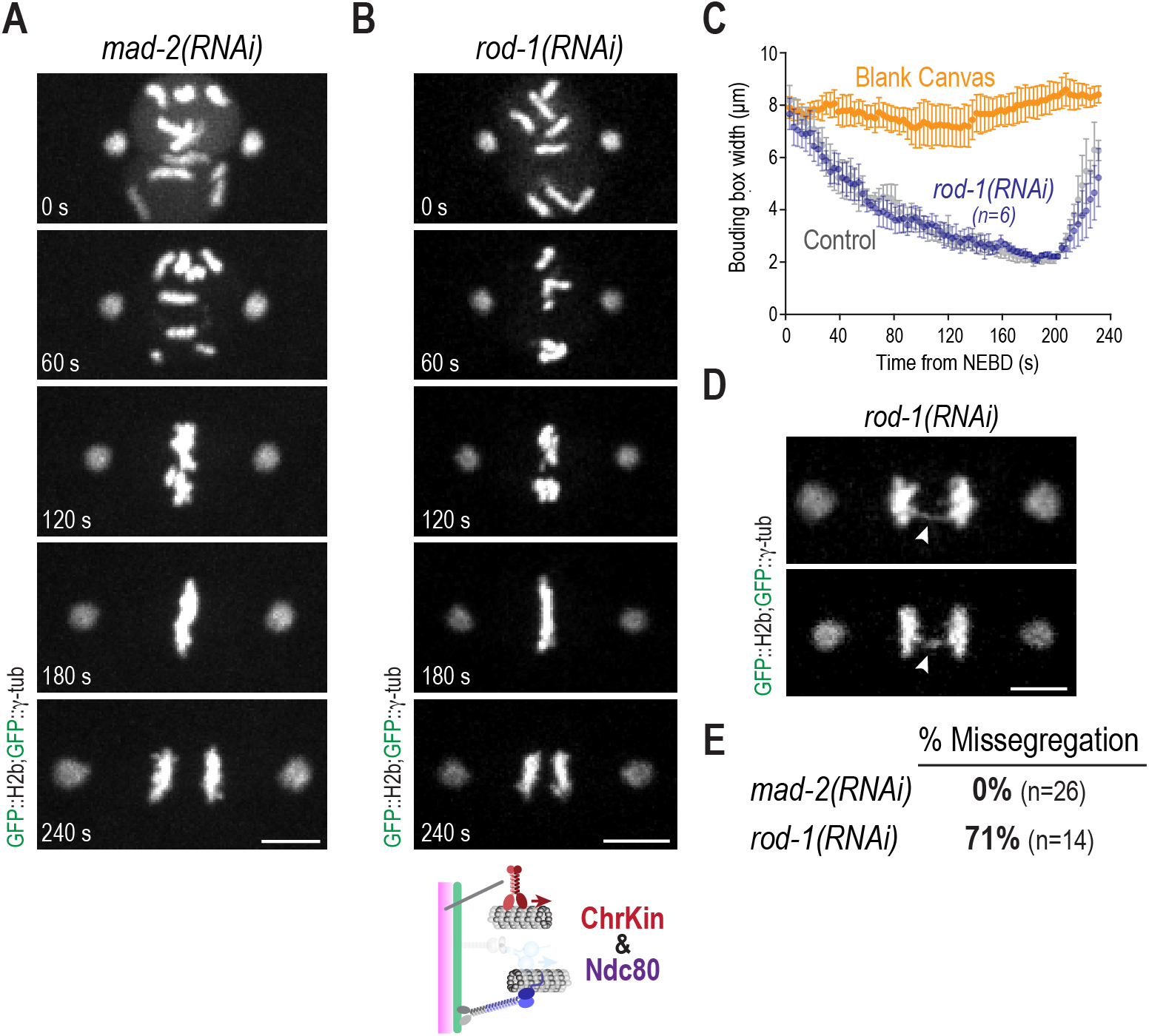
Analysis of checkpoint inhibition and of the chromokinesin–Ndc80 module combination created by removal of kinetochore dynein. **(A)** Chromosome dynamics in embryos depleted of the spindle checkpoint protein MAD-2. Loss of MAD-2 does not have any significant effect on chromosome segregation (*see also panel E*). Scale bar, 5 µm. **(B)** Chromosome dynamics in embryos lacking the kinetochore dynein module, where chromokinesin and the Ndc80 module are present. Scale bar, 5 µm. **(C)** Quantification of chromosome dispersion on the spindle performed as in Fig. 1D. The Control and Blank Canvas curves are the same as in Fig. 1D and are plotted to aid comparison. *n* is number of embryos analyzed. **(D)** Examples of anaphase segregation defects observed in the absence of the kinetochore dynein module. Scale bar, 5 µm. **(E)** Summary of missegregation events observed in anaphase of one-cell embryos. *n* is number of embryos imaged. Inhibition of the spindle checkpoint does not explain the segregation defect observed in the absence of the kinetochore dynein module.

**Figure S3.**
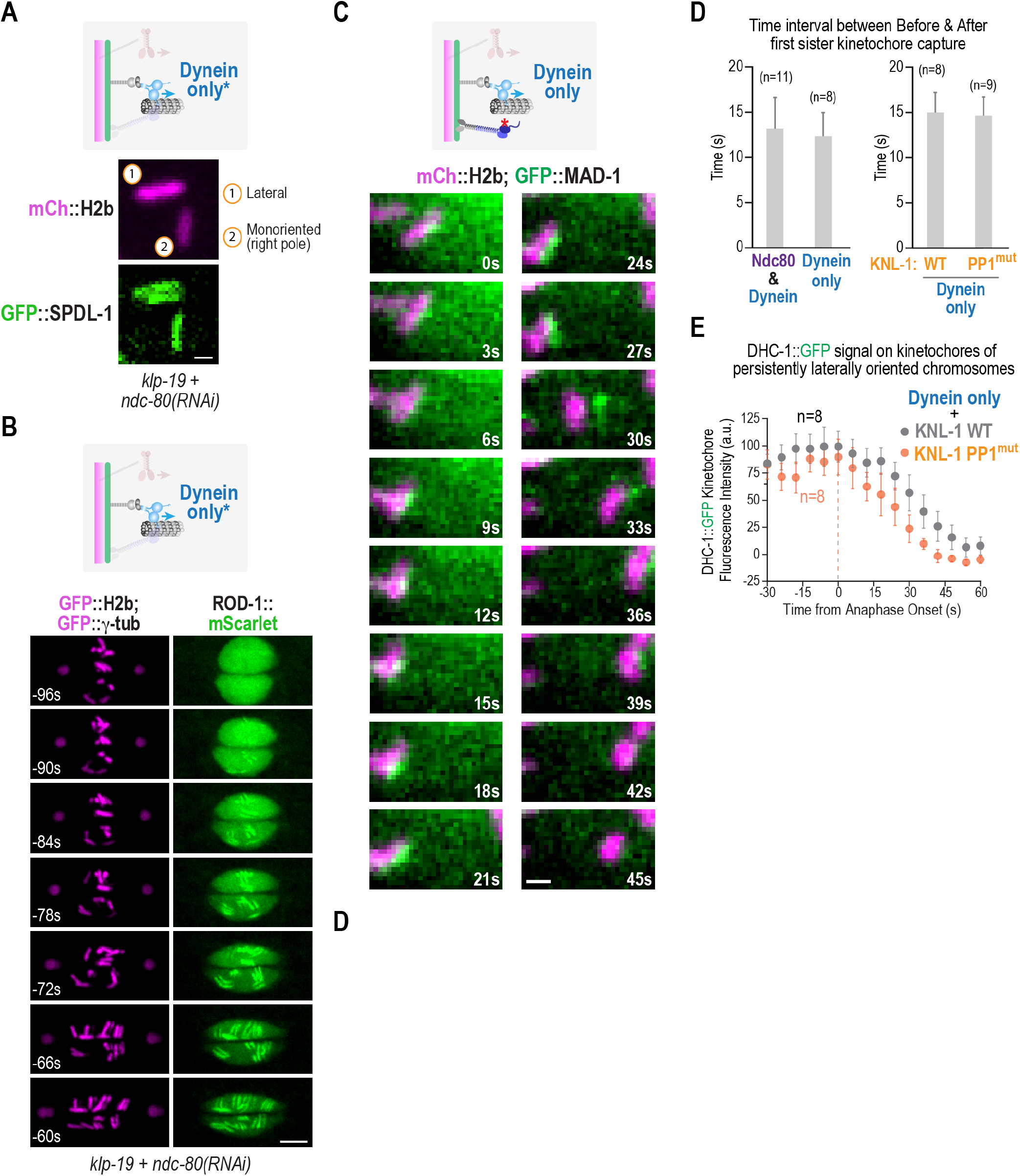
Imaging of GFP::SPDL-1, ROD-1::mScarlet and GFP::MAD-1, and supporting data for analysis of Ndc80 module and protein phosphatase 1 inhibitions. **(A) & (B)** Images of *in situ*-tagged GFP::SPDL-1 and ROD-1::mScarlet in *klp-19+ndc-80(RNAi)* embryos. As the mutant NDC-80 transgene was not present, this condition is labeled Dynein Only*. SPDL-1 behaved similarly to DHC-1, in that kinetochore-autonomous removal was observed. However, ROD-1 behaved distinctly–its levels were maintained at oriented kinetochores. Scale bars, 1 µm in *(A)* and 5 µm in *(B)*. **(C)** Image sequence of GFP::MAD-1 in the kinetochore dynein only state. The high signal of GFP::MAD-1 in the spindle region, together with its later recruitment, made imaging its dynamics on single kinetochores challenging. Nonetheless, chromosomes with clear kinetochore MAD-1 signal exhibited orientation-coupled removal from kinetochores. Scale bar, 1 µm. **(D)** Interval between the Before and After timepoints analyzed in Fig. 5A. The same interval is also plotted for the condition analyzed in Fig. 5B, although DHC-1 signal intensity before orientation was not measured. **(E)** Plot of DHC-1::GFP signal on persistently lateral chromosomes after anaphase onset for the indicated conditions. Error bars are the 95% confidence interval.

## METHODS

### C. elegans strains

*C*. *elegans* strains are described in *Table S1*. All strains were maintained using standard C. elegans growth media and imaged at 20°C. Endogenous locus GFP tagging was performed using CRISPR/Cas9^53^ at the *dhc-1* locus (dynein heavy chain) and the *klp-19* locus (chromokinesin). For both *dhc-1::gfp* and *gfp::klp-19* the repair template contained two homology arms, a linker sequence (GGRAGSG) and a sequence encoding GFP. GFP integrations were confirmed by PCR. For details on gRNAs, see *Table S2*.

### RNA-mediated interference

DNA templates were generated via PCR using the primers as specified in *Table S3*, and subsequently purified using a QIAquick PCR Purification Kit (Qiagen). Single-stranded RNA was generated from each DNA template using a MEGAscript^TM^ T3 and T7 Transcription Kit (Invitrogen), and subsequently purified using a MEGAclear^TM^ Transcription Clean-Up Kit (Invitrogen). Double-stranded RNA (dsRNA) was generated by annealing the single-stranded RNAs at 37°C for 30 minutes^43^. 36-46h before dissection and embryo imaging, the dsRNA was injected into L4 hermaphrodites, which were maintained at 20°C. All RNAi experiments were performed using 1 mg/ml individual dsRNAs, or 1:1 or 1:1:1 mixtures of 1 mg/ml individual dsRNAs.

### One-cell embryo fluorescence microscopy and image analysis

One-cell embryos were dissected from adult hermaphrodites in M9 buffer, placed onto a microscope slide containing a 2% agarose pad, and subsequently covered with a 22x22 mm high-precision cover glass (No. 1.5H, Marienfeld).

Embryos were imaged on a spinning-disk confocal (Revolution XD Confocal System; Andor Technology) with a confocal scanner unit (CSU-10, Yokogawa Corporation) attached to an inverted microscope body (TE2000-E, Nikon), illuminated using solid-state 100 mW lasers using either a 60X or 100X 1.4 NA Plan Apochromat oil objective (Nikon), and an EMCCD camera (iXon DV887, Andor Technology) (Desai Lab, San Diego).

One-cell embryos were also imaged on a spinning-disk confocal (CSU-W1 Confocal System, Nikon) with a confocal scanner unit (CSU-W1, Yokogawa Corporation) attached to an inverted microscope body (ECLIPSE Ti2-E, Nikon), illuminated using solid-state 200 mW lasers using either a 60X or 100X 1.4 HP Plan Apochromat oil objective (Nikon), and an sCMOS camera (Prime 95B, Teledyne Photometrics) (Cheerambathur Lab, Edinburgh).

For localization analysis of NDC-80::GFP, GFP::KLP-19 and DHC-1::GFP, 5 x 1.5 µm z-stacks were acquired every 10s and for KNL-1::GFP every 3s. For single chromosome localization analysis of DHC-1::GFP, 5 x 1.5 µm z-stacks were acquired every 3 s starting ∼1 min after NEBD and maximum intensity projections (MIPs) generated using Image J (Fiji). Subsequently, the fluorescent background was subtracted and a rectangular box (0.4 x 1.6 µm) fitted adjacent to the mCherry::H2B signal (chromosome) encapsulating kinetochore dynein to obtain the average DHC-1::GFP intensity.

For minimum bounding box (MMB) analysis, 5 x 1.5 µm z-stacks were acquired every 3 s, MIPs generated using Image J (Fiji) and rotated to position the spindle poles horizontally. Fluorescence intensity for all MIPs in the series was normalized, converted to 8-bit and the fluorescence background subtracted. Subsequently, to each MIP image a MMB was fitted (pixel value > 0, i.e. fluorescent signal from GFP::H2B) to measure chromosome positioning.

For chromosome orientation analysis, 5 x 1.5 µm z-stacks were acquired every 2 s for GFP::H2B or 3 s for DHC-1::GFP, MIPs generated using Image J (Fiji) and rotated to position the spindle poles horizontally. Subsequently, chromosome angles were determined by fitting a line along each chromosome axis and measuring the smallest angle between the chromosome axis and spindle pole-to-pole axis.

For EBP-2 imaging, a single z-section was acquired every 800 ms using 100 ms exposure, starting 1 min after NEBD. Kymographs were generated from a 5-pixel width line drawn pole-to-pole using KymographClear ^54^. Summed-intensity projection (SIP) images were generated by adding 38 EBP-2 frames starting 1 min after NEBD and subtracting the background signal. Intensity profiles were generated from these SIPs, by drawing a rectangular box (1.5 x 22 µm) encapsulating both spindle poles and averaging of the pixel intensities of each column within.

---

**Table S1:** C. elegans Strains.

| STRAIN DESCRIPTION | SOURCE | IDENTIFIER |
| --- | --- | --- |
| *C. elegans* N2 Bristol | Caenorhabditis Genetics Center | http://www.cg.c.cbs.umn.edu/strain.php?id=10570 |
| *unc-119(ed3) III; ruls32[pAZ132; pie-1/GFP::histone H2B] III; ddls6 [GFP::tbg-1; unc-119(+)] V* | Oegema et al. 2001, PMID: 11402065 | TH32 |
| *unc-119(ed3) III; ltIs37 [pAA64; pie-1/mCherry::his-58; unc-119 (+)] IV* |  | OD56 |
| *unc-119(ed3) III; ltSi120[[pDC170;Pndc-80:ndc-80 reencoded; cb-unc-119(+)]II #3* | Cheerambathur et al.,2013, PMID: 24231804 | OD611 |
| *unc-119(ed3) III; ltSi120[[pDC170;Pndc-80:ndc-80 reencoded; cb-unc-119(+)]II #3; ruls32[pAZ132; pie-1/GFP::histone H2B] III; ddls6 [GFP::tbg-1;unc-119(+)] V* | Cheerambathur et al.,2013, PMID: 24231804 | OD613 |
| *unc-119(ed3)III; ltSi129[pDC181;Pndc-80:NDC-80 (100,144,155AAA) reencoded; cb-unc-119(+)]II #2 ; ruls32[pAZ132; pie-1/GFP::histone H2B] III; ddls6 [GFP::tbg-1; unc-119(+)] V* | Cheerambathur et al.,2013, PMID: 24231804 | OD644 |
| *unc-119(ed3)III; ltSi122[pDC178;Pndc-80:NDC-80 (8,18,44,51AAAA) reencoded; cb-unc-119(+)]II #1; ruls32[pAZ132; pie-1/GFP::histone H2B] III; ddls6 [GFP::tbg-1; unc-119(+)] V* | Cheerambathur et al.,2013, PMID: 24231804 | OD688 |
| *unc-119(ed3)III; ltSi560 [pPLG014; Pmex-5::GFP::his-11::tbb-2_3'UTR, tbg-1::gfp::tbb-2_3'UTR; cb-unc-119(+)]V* | Kim et al., 2016,PMID: 26953348 | OD1702 |
| *unc-119(ed3)III; ltSi710[pDC267;Pndc-80:NDC-80 (66,96,100,125,144,155AAAAAA) reencoded; cb-unc-119(+)]II#1 ; ruls32[pAZ132; pie-1/GFP::histone H2B] III; ddls6 [GFP::tbg-1; unc-119(+)] V* | Cheerambathur et al.,2017, PMID: 28535376 | OD2312 |
| *dhc-1(lt45; dhc-1::gfp) I; unc-119(ed3) III?; ltIs37 [pAA64; pie-1/mCherry::his-58; unc-119 (+)] IV* | This Study | OD2956 |
| *lt53[knl-1::GFP::tev::loxP::3xFlag]III; ltIs37 [pAA64; pie-1/mCHERRY::his-58; unc-119 (+)] IV* | This Study | OD3075 |
| *lt53[knl-1::GFP::tev::loxP::3xFlag]III; ltSi711[pDC267;Pndc-80:NDC-80(66,96,100,125,144,155AAAAAA) reencoded; cb-unc-119(+)]II#1 ; ltIs37 [pAA64; pie-1/mCherry::his-58; unc-119 (+)] IV* | This Study | OD3083 |
| *lt53[knl-1::GFP::tev::loxP::3xFlag]III; ltSi129[pDC181;Pndc-80:NDC-80 (100,144,155AAA) reencoded; cb-unc-119(+)]II #2; ltIs37 [pAA64; pie-1/mCherry::his-58; unc-119 (+)] IV* | This Study | OD3121 |
| *klp-19(lt118[gfp::klp-19])III; ltIs3 [pAA64;pie1/mCherry::his-58; unc-119 (+)] IV* | This Study | OD3192 |
| *ndc-80(lt54[ndc-80::GFP::tev::loxP::3xFlag])IV; unc-119(ed3) III?; ltIs37 [pAA64; pie-1/mCherry::his-58; unc-119 (+)] IV* | This Study | OD3300 |
| *dhc-1(lt45; dhc-1::gfp) I; ltSi120[[pDC170;Pndc-80:ndc-80 reencoded; cb-unc-119(+)]II #3; unc-119(ed3) III?; ltIs37 [pAA64; pie-1/mCherry::his-58; unc-119 (+)] IV* | This Study | OD3630 |
| *dhc-1(lt45; dhc-1::gfp) I; ltSi711[pDC267;Pndc-80:NDC-80 (66,96,100,125,144,155AAAAAA) reencoded; cb-unc-119 (+) ]II#1; unc-119(ed3) III?; ltIs37 [pAA64; pie-1/mCherry::his-58; unc-119 (+)] IV* | This Study | OD3631 |
| *knl-1(lt53[knl-1::GFP::tev::loxP::3xFlag])III ltSi711 [pDC267;Pndc-80:NDC-80 (66,96,100,125,144,155AAAAAA) reencoded; cb-unc-119(+)]II#1; ltIs37 [pAA64; pie-1/mCHERRY::his-58; unc-119 (+)] IV* | This Study | OD3633 |
| *dha6[ebp-2::mNG:::tev::loxP::3xFlag]II* | This Study | DKC27 |
| *[ ItSi597[pDC202 ; Pknl-1::mCherry::knl-1::knl-1 3'UTR; cb-unc-119(+)] ; dhc-1 (It45[dhc-1::gfp]) ]I; ItSi711[pDC267;Pndc-80:NDC-80(66,96,100,125,144,155AAAAAA) reencoded; cb-unc-119(+)]II#1; unc-119(ed3)III? ; Itls37 [(pAA64) pie-1p::mCherry::his-58 + unc-119(+)] IV* | This Study | DKC133 |
| *[ dhaSi32[pDC646; Pknl-1::knl-1 reencoded (RRASA) :: mCherry::knl-13'UTR; cb-unc-119(+)]#1; dhc-1 (It45[dhc-1 ::gfp]) ]I; ItSi711[pDC267;Pndc-80:NDC-80 (66,96,100,125,144,155AAAAAA reencoded; cb-unc- 119(+)]II#1; unc-119(ed3)III? ; Itls37 [(pAA64) pie-1p ::mCherry::his-58 + unc-119(+)] IV* | This Study | DKC157 |
| *Itls37 [(pAA64) pie-1p::mCherry::his-58 + unc-119(+)] IV; (It39[gfp::tev::loxP::3xFlag::mdf-1])V)* | This Study | DKC231 |
| *ItSi120[pDC170;Pndc-80:NDC-80 reencoded; cb-unc-119(+)]II #3; unc-119(ed3)III? ; Itls37 [(pAA64) pie-1p::mCherry::his-58 + unc-119(+)] IV; (It39[gfp::tev::loxP::3xFlag::mdf-1])V)* | This Study | DKC232 |
| *ItSi711[pDC267;Pndc-80:NDC-80 (66,96,100,125,144,155AAAAAA) reencoded; cb-unc-119(+)]II#1; unc-119(ed3)III? ; Itls37 [(pAA64) pie-1p ::mCherry::his-58 + unc-119(+)] IV; (It39 [gfp::tev ::loxP :: 3xFlag::mdf-1])V)* | This Study | DKC233 |
| *dha6[ebp-2::mNG:::tev::loxP::3xFlag]II; knl-3 (dha19 [mscarlet-I^3XFLAG::knl-3] V* | This Study | DKC338 |
| *ruls32 [pie-1p::GFP::H2B + unc-119(+)] III. ddls6 [tbg-1::GFP + unc-119(+)] V.* | This Study | DKC393 |
| *ruls32 [pie-1p::GFP::H2B + unc-119(+)] III. ddls6 [tbg-1::GFP + unc-119(+); he279[Δebp-1, ΔY59A8B.25, Δebp-3] V;* | This Study | DKC394 |
| *spdl-1(dha113 (gfp::spdl-1) )II knl-1( It75[knl-1::mCherry]) III* | This Study | DKC594 |
| *rod-1(dha112 (ROD-1::mScarlet-I) )I; knl-1(It53[knl-1 ::GFP ::tev::loxP::3xFlag]))III* | This Study | DKC597 |
| *rod-1(dha112 (ROD-1::mScarlet-I) )I; ruls32 [pie-1p:: GFP::H2B + unc-119(+)] III. ddls6 [tbg-1::GFP + unc-119(+)] V.* | This Study | DKC760 |
| *spdl-1(dha113 (gfp::spdl-1) )II; unc-119(ed3) III; Itls37 [pAA64; pie-1/mCherry::his-58; unc-119 (+)] IV* | This Study | DKC761 |

**Table S2:**
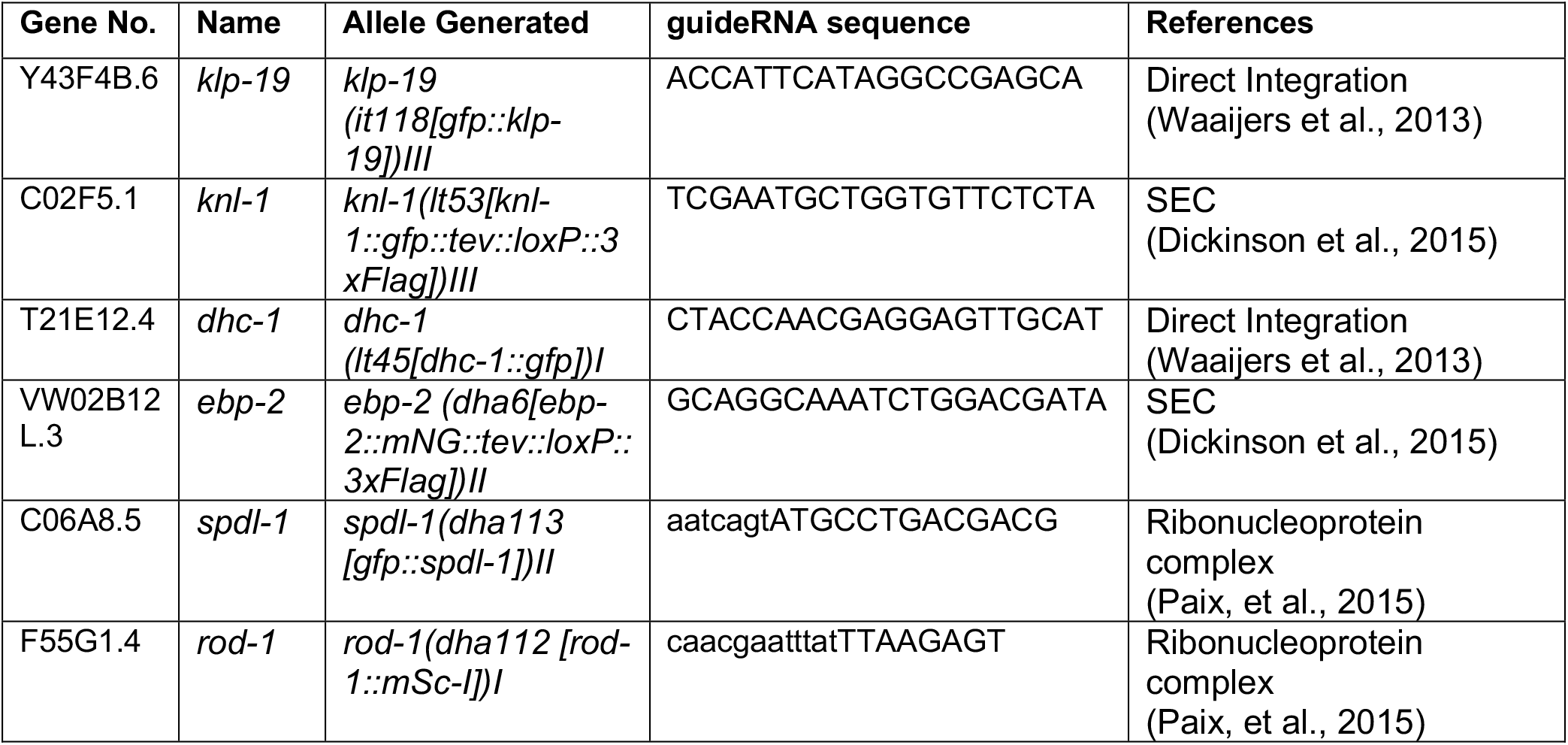
CRISPR gRNAs.

**Table S3:**
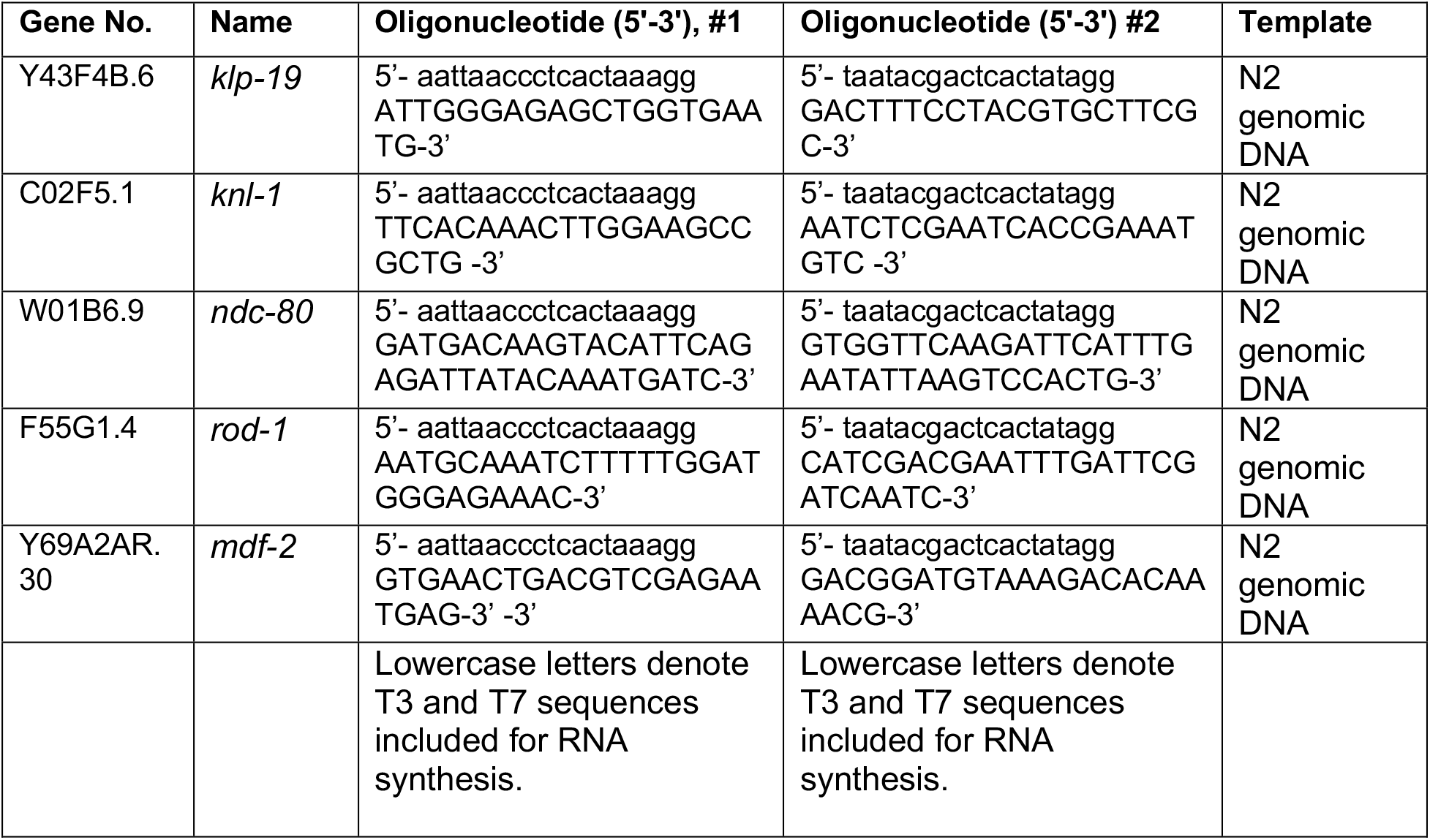
dsRNAs used in this study.

